# Investigating the role of long non-coding RNA in hypertrophic cardiomyopathy

**DOI:** 10.1101/2025.07.26.666851

**Authors:** Graham A. Branscom, Michael Morley, Jonathan J. Herrera, Jaime M. Yob, Sharlene M. Day

## Abstract

Long non-coding RNA (lncRNA) are transcripts that do not typically code for protein but have essential roles in the regulation of transcription and translation in health and disease. The objective of this study was to identify potential lncRNAs that could play a role in the pathophysiology of hypertrophic cardiomyopathy (HCM). We analyzed RNA-Seq data for lncRNA expression from a mouse model of HCM and cross-referenced transcripts to a published human HCM tissue dataset. We identified a total of 9,140 lncRNA transcripts in the mouse dataset, of which 35 were differentially expressed between transgenic *TNNT2* Δ160 mice (TG) and non-transgenic mice (nTG, p-*adj* < 0.05). Of these, 13 had a human ortholog as predicted by ortho2align. We used the computational tools MiPepid, AlphaFold, and PhyloCSF to predict potential micropeptides that could be coded for by these 13 mouse lncRNAs. We found that predicted micropeptides from 3 of these lncRNAs–*G730003C15Rik*, 9830004L10Rik, and Gm45012–have higher AlphaFold folding confidence metrics than random peptides or truly non-coding lncRNA negative controls (*p* < 0.05). Another 2 of these lncRNAs, *6330403L08Rik* and *2900072N19Rik*, have positive PhyloCSF scores, also indicating micropeptide coding potential. In summary, we developed a computational workflow to identify differentially expressed lncRNAs in a mouse model of HCM that can be prioritized for future experimental studies based on their cross-species conservation and micropeptide coding potential.

**NEW & NOTEWORTHY:** This is the first analysis of RNA-Seq data for lncRNA expression in an HCM mouse model and the first cross-species analysis of HCM lncRNA RNA-Seq data. Additionally, this study demonstrated a novel computational pipeline that combines several tools–RNA-Seq, MiPepid, AlphaFold, and PhyloCSF–to identify potential lncRNAs of interest from RNA-Seq data.

## INTRODUCTION

Long non-coding RNAs (lncRNAs) are transcripts greater than 200 nucleotides long that are typically annotated as not coding for protein.^1^ However, lncRNAs have a diverse array of biological roles, including transcriptional regulation of target genes, chromatin remodeling, and nuclear organization, among many others.^2,3^ Some lncRNAs are subject to similar transcriptional processing as mRNA, such as addition of a 5’ cap, polyadenylation, and splicing. Although lncRNAs were traditionally believed to all be non-coding, many lncRNAs have since been identified to contain small open reading frames (sORFs) that encode for micropeptides less than 100 amino acids long.^4-6^

Hypertrophic cardiomyopathy (HCM) is an inherited cardiovascular disorder marked by left ventricular (LV) hypertrophy, diastolic dysfunction, and fibrosis.^7^ Currently, several lncRNAs have been experimentally linked to cardiac hypertrophy.^8,9^ For instance, the lncRNA *Mhrt* is protective against induced pathological hypertrophy by binding to and inhibiting the enzyme Brg1.^10,11^ Other lncRNAs like Plscr4 regulate pathological hypertrophy by acting as molecular sponges for miRNAs, preventing their binding to downstream mRNA transcript targets.^12^

While some lncRNAs have been identified as dysregulated in HCM through methods like RT-qPCR,^13^ unbiased high-throughput techniques like total RNA-Seq and lncRNA-based microarrays have identified key lncRNAs that are differentially expressed (DE) in HCM.^14,15^ GSE130036 is the primary human HCM RNA-Seq dataset that includes lncRNA.^14^ These authors identified 150 DE lncRNAs with an absolute fold change (FC) > 2 and a q-value < 0.05, of which 67 were upregulated and 83 downregulated in HCM. Since this initial study was published, several other studies have conducted similar DE analyses of the lncRNA genes in this dataset and constructed predicted lncRNA-miRNA-mRNA networks that could regulate HCM.^16-18^ Other studies have found gene ontology (GO) terms that are significantly enriched among the DE genes in this dataset, yet none found terms related to non-coding RNA processing.^19,20^ In addition to traditional DE analysis of lncRNAs in HCM, over 100 lncRNA sORF-encoded micropeptides have been detected in primary cardiomyocytes exposed to hypertrophic agonists using Ribo-Seq which sequences ribosome-protected fragments.^21^

Typically, hundreds to thousands of lncRNAs are identified as being differentially expressed in disease compared to control samples.^14,15^ It is challenging to pare these down to a reasonable number of lncRNA candidates that can be prioritized for experimental studies. Available computational tools use crossspecies conservation metrics to identify evolutionary conserved regions in lncRNAs,^22^ which are more likely to serve essential functions in a particular organ. However, non-coding regions of the genome are typically less conserved than protein-coding regions due to weaker evolutionary constraints.^23,24^ While there are an estimated 30,000 to 60,000 lncRNA genes expressed in human, only 1,731 are computationally predicted to be conserved across human and mouse.^25-27^

This study aims to identify lncRNAs that may be important in the pathophysiology of HCM. We analyze an RNA-Seq data from a mouse model of HCM (*TNNT2* Δ160) and cross reference our findings to a human HCM dataset. After we identified lncRNAs that are DE in mice and possess a human ortholog, we employed various computational tools to identify micropeptide coding potential among these lncRNAs. These lncRNAs can be prioritized for future experimental investigation.

## MATERIALS AND METHODS

The overall methodological workflow is shown in Fig. 1A.

**Figure 1.**
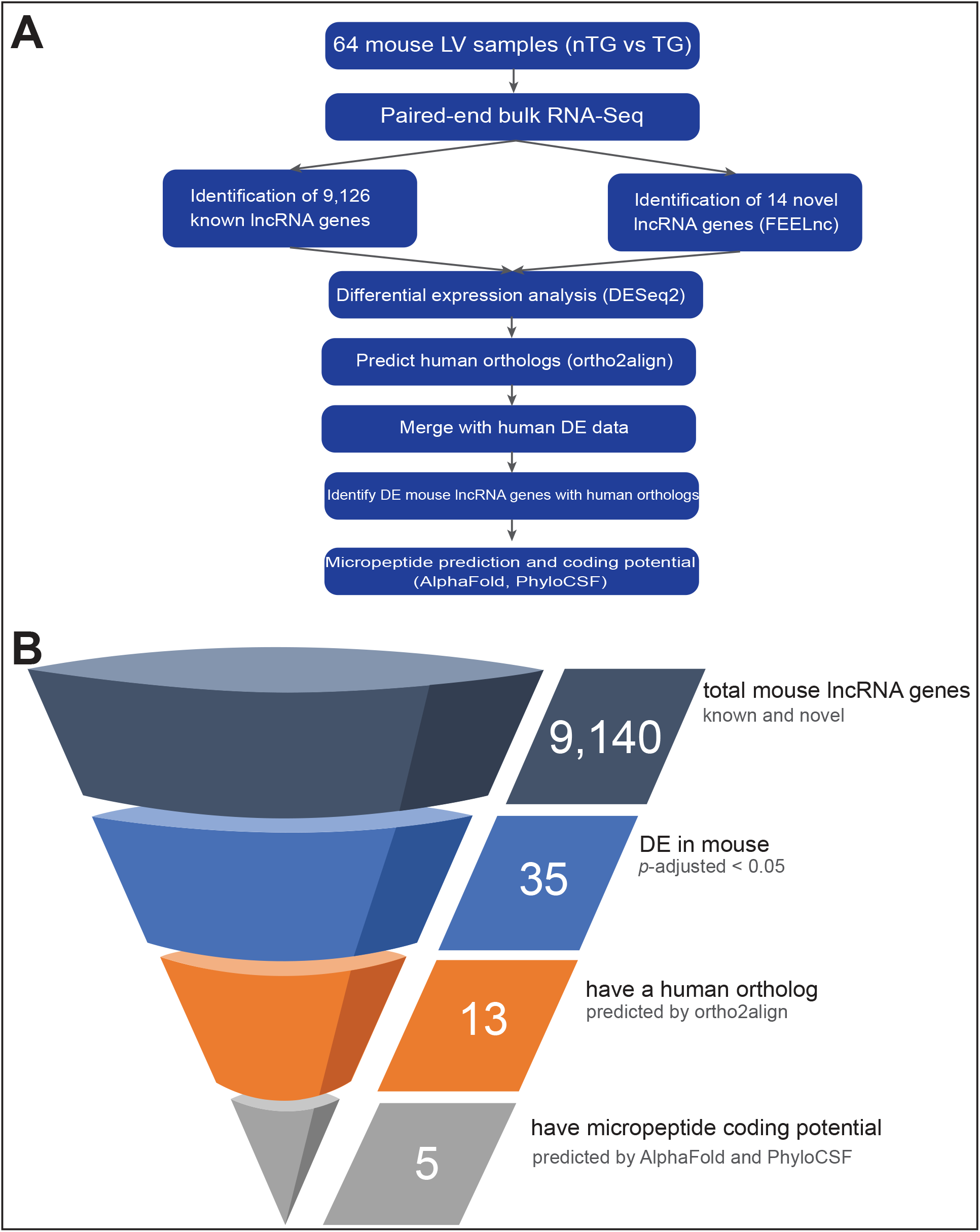
Scientific methodology A. The methodology workflow for this study. Mouse RNA-Seq data includes left ventricular tissue from 64 mice (n=31 transgenic *TNNT2* Δ160 (TG), n=33 non-transgenic (nTG)). Within TG, there are 17 males and 14 females. Within nTG, there are 15 males and 18 females. Within the TG group, there were 14 HIIT and 17 Sed mice, and 18 HIIT and 15 Sed within the nTG group. Human RNA-Seq data includes myocardial tissue from 37 patients (n=28 HCM, n=9 non-failing). Among HCM samples, 19 were from males and 9 were from females. Among NF samples, 8 were from males and 1 was from a female. B. The filtering of genes used when analyzing the mouse RNA-Seq data. *p*-values were corrected under the Benjamini-Hochberg procedure. TG: transgenic, nTG: non-transgenic, *p*-adj: adjusted p-value, DE: differentially expressed.

### Mouse model

C57BL/6J mice (n=64) were obtained from Dr. Jil Tardiff’s laboratory.^28,29^ This included 33 non-transgenic (nTG) and 31 transgenic (TG) mice which had a deletion of Glu160 (Δ160E) in *TNNT2* driven by the α-MHC promoter. Mutant cTnT replaced 70% of endogenous cTnT. This model recapitulates key features of human HCM including diastolic dysfunction and hypercontractility. There were 17 TG males and 14 TG females and 15 nTG males and 18 nTG females. In a concurrent study, mice were run under a high intensity interval training (HIIT) and sedentary (Sed) protocol (unpublished). Within the TG group, there were 14 HIIT and 17 Sed mice, and 18 HIIT and 15 Sed within the nTG group. The HIIT group was run on a treadmill with high intensity intervals for a total of 30 minutes, 3 days per week, for 6 weeks. We investigated overall differences in TG vs nTG mice with exercise and sex as covariates.

### RNA-Seq data collection

Total RNA was extracted from the LVs of mice using random hexamer primers. RNA was subject to ribosomal depletion and bulk and paired RNA-Seq reaching a high depth. The dataset is made publicly available under GSE297707.

Human RNA-Seq data was downloaded from a published dataset (GSE130036). In this study, myocardial tissue was obtained from 37 patients (28 patients with HCM and 9 non-failing (NF) donor controls). There were 19 men and 9 women within HCM, 8 NF men, and 1 NF woman. This RNA was subject to total RNA extraction and strand-specific library preparation.^14^

### Differential expression analysis

lncRNA transcripts were identified using a full bioinformatics pipeline (Fig. 1A). The FASTQ files were first subject to <u>FASTQC</u> (v0.12.0) and <u>MultiQC</u> (v1.23) to assess read quality. The adapter sequences were cut using <u>Cutadapt</u> (v4.9) and then aligned to the GENCODE primary assembly mouse genome (GRCm38/mm10) with <u>STAR</u> (v2.7.11b).^30^ <u>Cufflinks</u> and Cuffmerge (v2.2.1) were used for transcript assembly.^31^ The Cuffmerge output GTF file was compared with 60,411 known lncRNA genes from GENCODE, which found 9,126 known unique lncRNA genes across all our mouse samples. <u>FEELnc</u> (v0.2.1) found 14 novel lncRNA genes,^32^ for a total of 9,140 lncRNA genes. <u>RSEM</u> (v1.3.3) was used to generate count matrices.^33^ These matrices were imported into R (v4.4.2) using tximport and analyzed with DESeq2 in R via RStudio (v2024.12.1).^34,35^ We used <u>ortho2align</u> (v1.0.5) to find known and predicted human orthologs of mouse lncRNA genes.^36^

Human RNA-Seq abundance matrices were downloaded from GSE130036. We used the author’s published spreadsheet of genes identified by RNA-Seq to identify lncRNA genes and analyzed the data using DESeq2.^14,35^

A variance stabilization transformation was used on the data for count heatmaps and principal component analyses (PCA). DESeq2 analysis used independent filtering with Benjamini-Hochberg (BH)-corrected *p*-values (*p*-adj) with an alpha of 0.05 and a FC of one. Genes were DE if they had a *p*-adj < 0.05, regardless of their FC value since FC values were very low (mean log_2_FC among mouse genes with *p*-adj < 0.05 was -0.35 (SD 0.73); -0.228 (SD 1.23) for human). Other R packages used included pheatmap, EnhancedVolcano, ggVennDiagram, tidyverse, and dplyr. All computation except the analysis in R was done on the Penn Medicine Academic Computing Services High Performance Computing cluster. Code and other resources can be found in Supplementary Code C1.

We searched each DE lncRNA on the RNAcentral database for known tissue-specific expression data.^37,38^

### Gene ontology analysis

GO analysis was run on both mouse and human RNA-Seq data using all ontologies (biological process, molecular function, and cellular component). For over representation/enrichment analysis (ORA), we input only genes (protein-coding and non-coding) with a *p*-adj < 0.05 (n=1,052 for mouse, n=7,630 for human). For gene set enrichment analysis (GSEA), we input all genes (protein-coding and non-coding) regardless of *p*-adj (n=16,892, n=16,741 for mouse and human, respectively). We used the R packages clusterProfiler and enrichPlot. Enriched terms had a BH-adjusted *p*-values < 0.05.

### Computational prediction of micropeptide coding potential

13 lncRNA genes that had *p*-adjusted < 0.05 and possessed human orthologs were assessed for micropeptide coding potential (Fig. 1B). The contiguous exon DNA sequence of their canonical isoforms was run through <u>MiPepid</u>, a machine-learning based tool for identifying sORFs that may encode for micropeptides.^38^ These sORFs were translated using EMBOSS <u>transeq</u> into micropeptides and then run through AlphaFold2 via the ColabFold interface (v1.5.5).^40,41^ We used mean predicted local distance difference test and predicted aligned error to assess coding potential. We compared these predictions to micropeptides coded by sORFs from true non-coding (TNC) lncRNAs, false non-coding (FNC) lncRNA (i.e. lncRNAs that encode micropeptides), and randomly generated peptide sequences. To find TNC and FNC sequences, the entire mouse lncRNA transcriptome (n=155,417 transcripts) was run through MiPepid. The TNC and FNC sORFs were the top 120 sORFs with the highest non-coding and coding probabilities from the MiPepid output, respectively. For the randomly generated micropeptides, we generated 120 peptides using the Python (v3.9.19) random module. We stratified the results by 10 amino acid length increments since sORF length is known to be a strong predictor of micropeptide coding potential by MiPepid.^39^ Independent two sample t-tests compared scoring differences between micropeptide types. AlphaFold structures were rendered via UCSF ChimeraX (v1.9.0).^42^ The number of predicted sORFs for each gene ran through AlphaFold was compared to the number for all lncRNA genes in mouse, and significance was determined with a permutation test.

These 13 lncRNA genes were also run through PhyloCSF, a tool that uses a multi-species alignment to identify protein-coding regions among lncRNA sequences.^22^ We employed the methodology described in a previous study to load PhyloCSF as a track hub on the UCSC genome browser and analyzed the smoothed, approximate coding regions, candidate coding regions, and relative branch lengths tracks.^43^ The power track is a score from 0 to 1, with higher scores representing higher statistical confidence.

Figures were generated with ggplot in R and Plotly in Python and edited with Adobe Illustrator (v29.5.1).

## RESULTS

### Mouse lncRNA DE analysis

Using the computational pipeline outlined in Fig. 1A, we identified 9,140 unique lncRNA genes in mice. There were 77.87E6 (SD 9.45E6) reads per sample. A heatmap of the top 40 lncRNA genes shows that there is sample clustering primarily driven by genotype (Fig. 2A). PCA confirms that there is clustering present when samples are colored by their genotype (Fig. 2B), but clustering is not present when colored by sex (Fig. 2C). The outlier in the upper-left corner of both PCA plots had the highest percentage of unmapped reads among all samples (8.6%, average among all samples: 5.8%). TG vs nTG analysis resulted in the most DE genes (n=25) (Fig. 2D) compared to sex (n=4) (Fig. 2E) or exercise group (n=0). However, we also included analyses stratified by sex and exercise group (Supplemental Fig. S1) to minimize covariate effects and maximize the number of DE genes captured. 35 unique lncRNA genes were found among all the TG vs nTG comparisons. 11 were found only in TG vs nTG, 8 among TG HIIT vs nTG HIIT, 1 among TG males vs nTG males, and 15 were found across multiple comparison groups (Fig. 2F, Supplemental Fig. S1). Among these 35, 8 are upregulated in TG and 27 are downregulated in TG.

**Figure 2.**
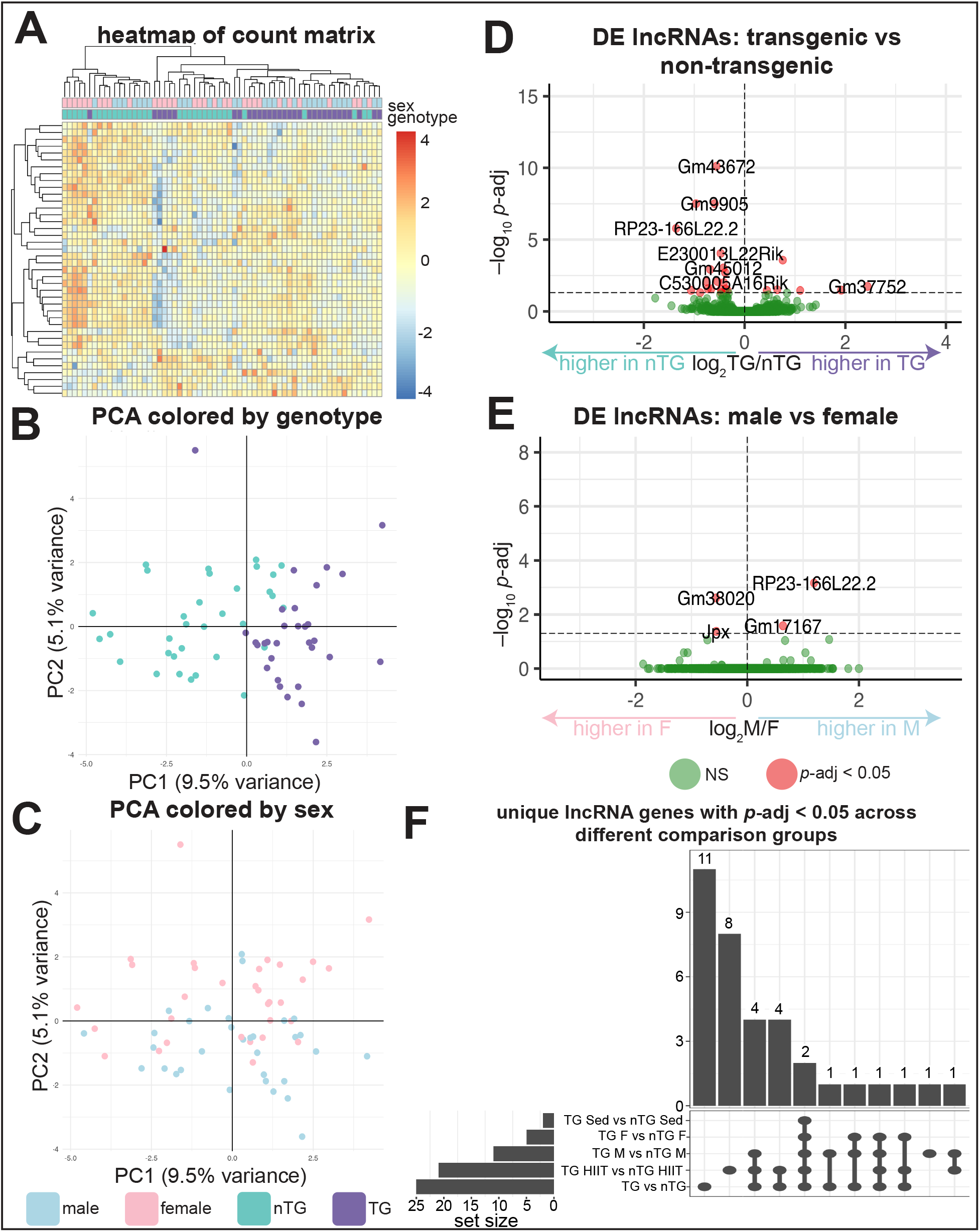
DE analysis of mouse RNA-Seq data A. Heatmap of the 40 lncRNA genes with the highest counts, annotated by genotype and sex. Both rows (genes, n=40) and columns (samples, n=64) were clustered. Count data was subject to a variance stabilizing transformation prior to clustering and scaled gene-wise. B. PCA of the top 500 genes with the highest variance after variance stabilizing transformation, colored by genotype. C. Same PCA as panel B but colored by sex. D. Volcano plot from the DE analysis of all the lncRNA genes for the *TNNT2* Δ160 transgenic (TG) vs nontransgenic (nTG) comparison. *y*-axis denotes *p*-values adjusted with the Benjamini-Hochberg procedure. The horizontal line above the x-axis indicates a *p*-adjusted cutoff of 0.05. E. Similar to panel D but DE for male vs female. F. Bar chart depicting the number of unique lncRNAs that have *p*-adjusted < 0.05 across five different comparisons: TG vs nTG (among all samples, n=64), TG M vs nTG M (among males only, n=32), TG F vs nTG F (among females only, n=32), TG HIIT vs nTG HIIT (among HIIT mice only, n=32), TG Sed vs nTG Sed (among sedentary mice only, n=32). Sex and exercise were treated as covariates. The bottom panel indicates which comparisons are included in each bar. PCA: principal component analysis, DE: differential expression, TG: transgenic, nTG: non-transgenic, M: male, F: female, HIIT: high intensity interval training, Sed: sedentary

These 35 genes in total, along with their human orthologs (if any) and known functions are in Supplemental Table T1. Only four of these genes have a known role. Two are known to be DE following irradiation,^44-46^ three are DE in cancer,^47-50^ and one called *ANRIL* regulates myocardial infarction and coronary artery disease.^51-53^ A screen of the RNAcentral database showed that 14 of the 35 lncRNAs have known tissue-specific expression, with 9 of these having known expression in the heart.

Over-representation analysis (ORA) analysis among all DE genes, both protein-coding and non-coding (n=1,052), revealed that “regulation of non-coding RNA (ncRNA) transcription” was an enriched term in the mouse dataset.

### Human lncRNA DE analysis

Previous studies have already conducted DE analysis on a human HCM dataset specifically for lncRNA genes.^14,16,18^ We reran the DE analysis to harmonize criteria with the mouse dataset (*p*-adj < 0.05) to enable valid comparisons. We found that of the 2,398 total unique lncRNA genes captured by RNA-Seq, 340 were DE (*p*-adj < 0.05) across all HCM vs NF samples and the sex-stratified comparisons. Specifically, 64 were DE among all HCM vs NF, 31 among HCM vs NF males, 24 among HCM vs NF females, and 221 were DE across multiple comparison groups. Of these 340 total genes, 139 were higher in HCM than NF and 198 were lower in HCM than NF. (Three have conflicting results, e.g. higher in all HCM vs all NF, but lower in HCM vs NF males).

ORA among all DE genes, both protein-coding and non-coding (n=7,630), revealed that “ncRNA processing,” “ncRNA 5’-end processing” and “ncRNA transcription” were enriched in the human dataset (*p*-adj < 0.05). Additionally, gene set enrichment analysis (GSEA) revealed that there was one term (“ncRNA processing”) that had a significant enrichment score (*p*-adj < 0.05). The running enrichment score, a measure of the cumulative enrichment, for ncRNA processing was higher for coding and non-coding genes that had both higher (higher among HCM) and lower (higher among NF) log_2_FC values, with a slightly skewed enrichment toward NF. This indicates that “ncRNA processing” consists of genes that are both upregulated and downregulated in HCM, with a slight bias toward downregulation (Fig. 3A).

**Figure 3.**
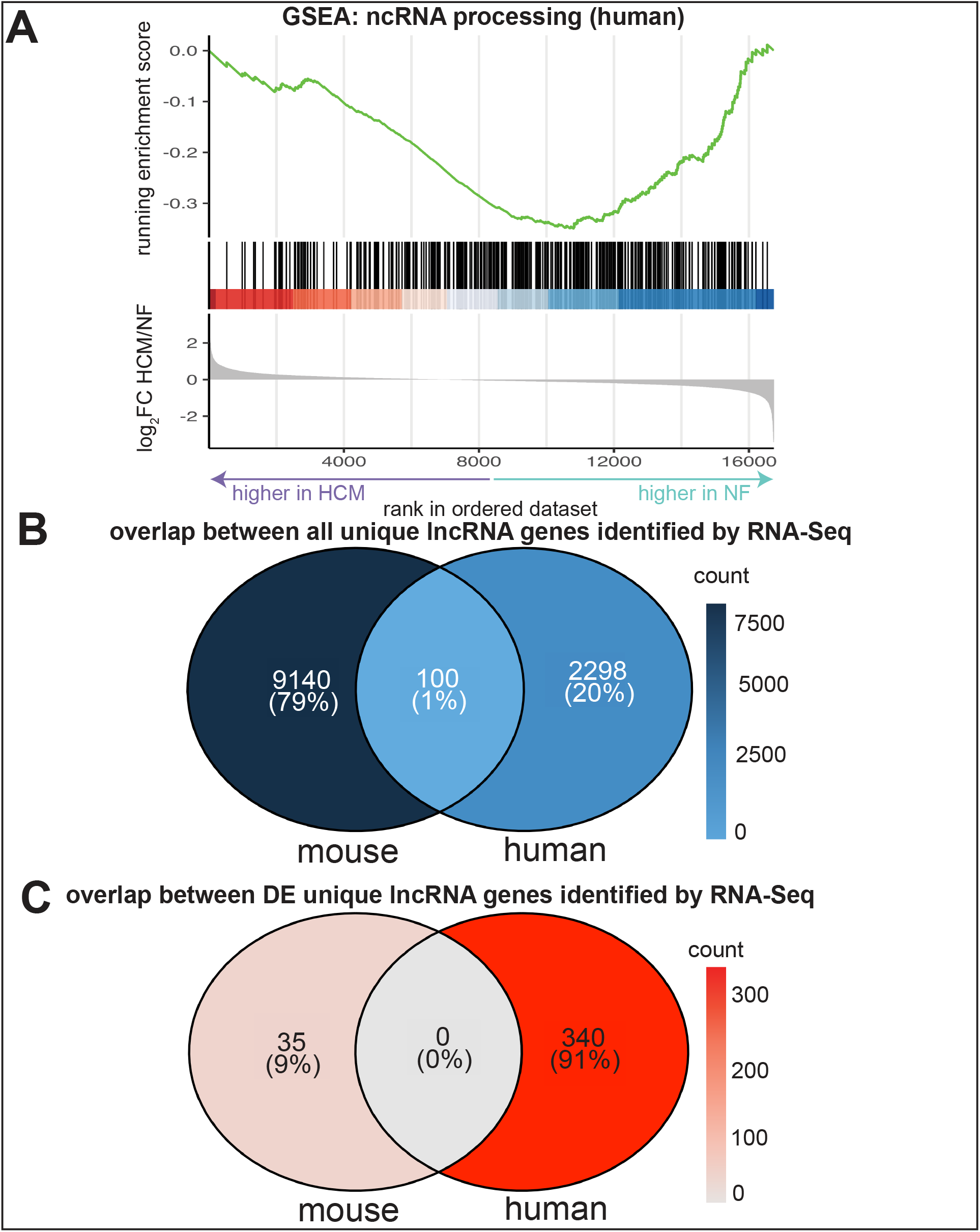
GO analysis and comparison between mouse and human RNA-Seq datasets A. Gene set enrichment analysis (GSEA) gene ontology (GO) analysis among all genes identified in the human dataset, both coding and non-coding (n=16,741) total. The non-coding RNA (ncRNA) processing GSEA term had a *p*-adjusted < 0.05. The *x*-axis denotes the rank of all the genes in the dataset using descending log_2_FC, from higher in HCM (left/lower rank) to higher in non-failing samples (right/higher rank). B. Overlap between all unique lncRNA genes identified by RNA-Seq in both the mouse and human datasets. No filtering was applied. C. Overlap between all unique lncRNA genes among the mouse and human datasets, but only for genes with a *p*-adjusted < 0.05, regardless of log_2_FC value. *P*-values are adjusted by the Benjamini-Hochberg procedure. GO: gene ontology, GSEA: gene set enrichment analysis, FC: fold change, HCM: hypertrophic cardiomyopathy, NF: non-failing

### Cross-species lncRNA comparison

Among all the lncRNA genes identified by RNA-Seq in mouse (n=9,140) and human (n=2,398), only 100 (0.87%) were shared across both species (Fig. 3B). Among the DE genes (*p*-adj < 0.05) in mouse (n=35) and human (n=340), 0 were shared across both species (Fig. 3C). Even using an unadjusted *p*-value < 0.05 as the DE criterion instead of the BH-adjusted *p*-value, there were still no shared DE genes between mouse and human.

### Micropeptide coding prediction

We ran micropeptide coding prediction analysis on the 13 lncRNA genes that were DE in mouse and had human orthologs. We chose these genes because they are potentially involved with HCM pathophysiology and are evolutionarily conserved. We first used MiPepid to predict small open reading frames (sORFs) in the lncRNA gene sequences. 9/13 of these transcripts had a significantly greater number of small open reading frames (sORFs) than the average of all 155,417 mouse lncRNA transcripts, making them more likely to encode for micropeptides (*p* < 0.05, Supplemental Fig. S2A). We then ran the peptide sequences coded by these predicted sORFs using AlphaFold. We followed a similar methodology to published studies that used the mean per-residue predicted local distance difference test (pLDDT) value from AlphaFold, which measures folding confidence, as a proxy for micropeptide coding potential.^54,55^ We found that the lncRNA *G730003C15Rik* has predicted micropeptides in the 1-10 amino acid length range that have higher mean pLDDT scores than the randomly generated peptides negative control, indicating greater confidence in these being true micropeptides (*p* < 0.01, Fig. 4A). *9830004L10Rik* and *Gm45012* have micropeptides within two and one different length ranges, respectively, with higher mean pLDDT scores than random peptides (p < 0.05, Fig. 4A).

**Figure 4.**
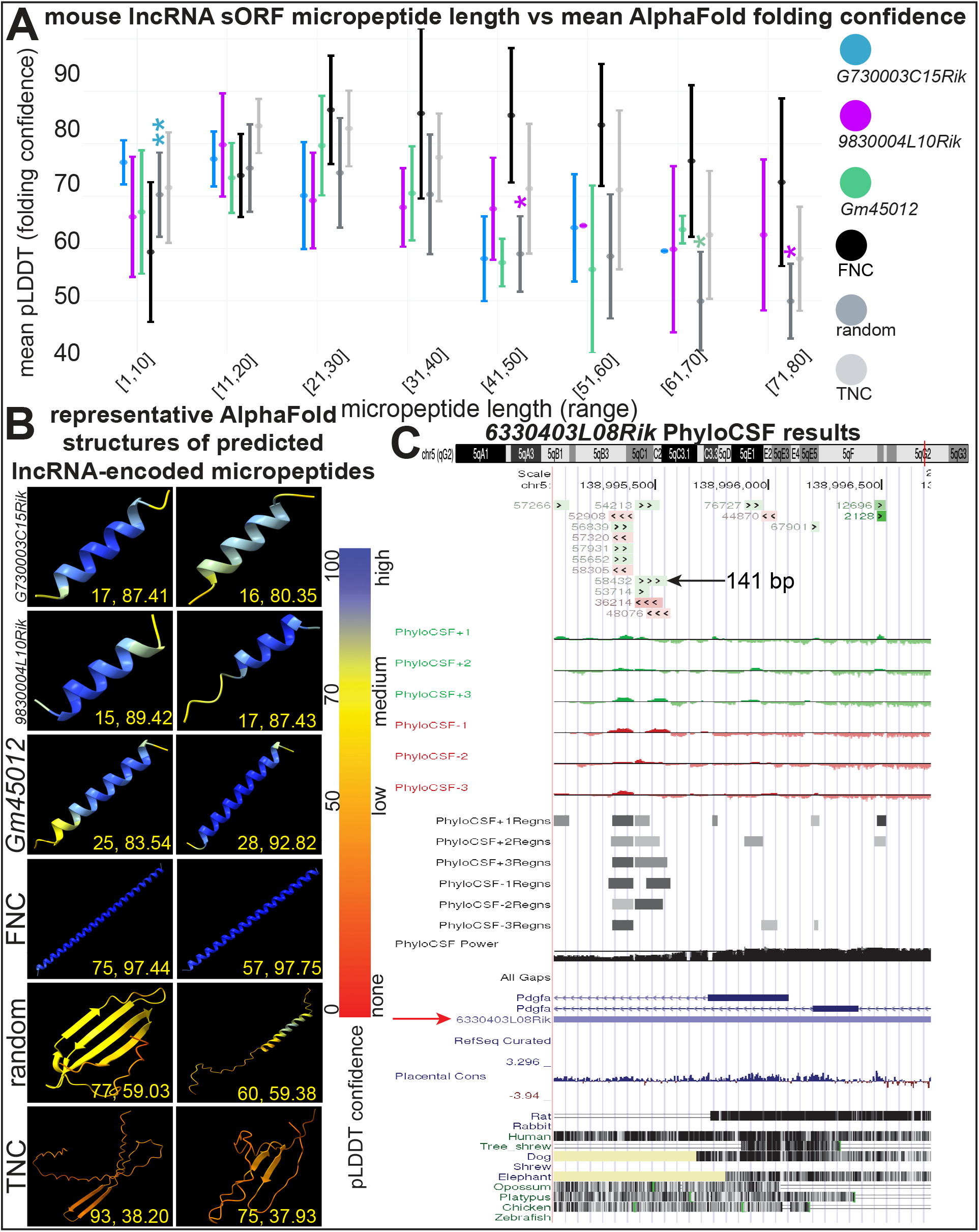
Predicting micropeptide coding potential A. Measures of folding confidence for micropeptides coded by the 13 DE (*p*-adj < 0.05) mouse lncRNAs with human orthologs, stratified by micropeptide length. Only lncRNA genes with significant results are shown. Mean per-residue predicted local distance difference test (pLDDT) scores are for micropeptides falling into buckets of length 10 amino acids long from 1 to 80. Higher pLDDT scores indicate lower folding confidence and higher micropeptide coding potential. There are n=70 micropetides for *G730003C15Rik*, n=233 for *9830004L10Rik*, and n=106 for *Gm45012*. There are 120 micropeptides each for false non-coding (FNC, i.e. true coding), random, and true non-coding (TNC), 15 for each length range. Error bars depict standard deviation. *p*-values were determined with independent two sample t-tests. *p*-values are denoted by asterisks sitting above and having the color of the two groups that they are comparing within a given length range subset. For simplicity, *p*-values are only shown when any of the three lncRNAs have a greater pLDDT than random or TNC micropeptides. B. Representative predicted micropeptides structures outputted by AlphaFold and rendered with UCSF ChimeraX. Two micropeptide-encoded sORFs from the 10 with the highest pLDDT scores are shown for *G730003C15Rik, 9830004L10Rik, Gm45012*, and FNC. Two from the 10 median scoring sORFs are shown for the random micropeptides. Two from the 10 with the lowest pLDDT scores are shown for TNC. Numbers in each box are listed as: micropeptide length (number of amino acids), pLDDT. Each residue is colored based on its pLDDT value according to the color gradient. C. PhyloCSF tracks in the UCSC Genome Browser for the mouse lncRNA *6330403L08Rik* in genome build GRCm38/mm10. The topmost track is the PhyloCSF novel track, indicating novel coding regions (green is on plus strand, red is on minus strand). Below that is the Smoothed PhyloCSF track for each of the six possible reading frames, which depicts the scores given by PhyloCSF’s hidden Markov model. Next is the PhyloCSF regions track for all reading frames, indicating the Smoothed track’s predicted coding regions. Next is the PhyloCSF Power track, which indicates a score from 0 to 1 indicating statistical confidence, with a higher score indicating higher confidence. Next are the genes located within this region. *6330403L08Rik* is marked with a red arrow. Lastly, there is the Conservation track of 100 vertebrates, including mouse, for reference. * *p* < 0.05, ** *p* < 0.01 sORF: small open reading frame, pLDDT: predicted local distance difference test, FNC: false non-coding (i.e. true coding), TNC: truly non-coding

We used another AlphaFold metric called predicted aligned error (PAE), where lower scores indicate higher confidence in micropeptide folding. We found that at the 1-10 range, *G730003C15Rik* had a lower mean PAE than both the random micropeptides and micropeptides from truly non-coding (TNC) lncRNAs, which served as another negative control (*p* < 0.05). At the 11-20 amino acids range, *9830004L10Rik* had a lower mean PAE value than the random peptides (*p* < 0.05, Supplemental Fig. S2B). Representative renderings of predicted micropeptides from the three lncRNAs with significant pLDDT and PAE results are shown in Fig. 4B. Higher scoring predicted micropeptides like those shown for the three lncRNA genes and the falsely non-coding (FNC) lncRNAs positive control tend to be alpha helices, whereas those for lower scoring micropeptides like TNC and random tended to be beta sheets or peptides with little secondary structure.

Two of the 13 mouse lncRNAs had positive PhyloCSF scores. PhyloCSF predicted that *6330403L08Rik* had 17 predicted sORF, with the longest being 141 base pairs long. Although that sORF had a low power score (0.255), five shorter sORFs had high power scores (0.598-0.956). Another lncRNA, *2900072N19Rik*, had one small nine base pair sORF with a positive smoother PhyloCSF score in 2/6 reading frames (Supplemental Fig. S3).

## DISCUSSION

In summary, we found 35 mouse lncRNAs that are DE in an HCM mouse model. 13 of these genes have a human ortholog, indicating that they are evolutionarily conserved.^24,56^ We found that 5/13 (38.5%) of these lncRNAs have predicted micropeptides with significant AlphaFold folding confidence metrics or PhyloCSF results. By conducting GO analysis among all DE mouse and human genes, we also found that there is one enriched term related to lncRNA processing in mouse and three in human, consisting of genes that are both upregulated and downregulated in the HCM model.

Although studies have analyzed RNA-Seq data for lncRNA expression among human HCM samples,^14^ a cross-species comparator study has not been previously reported. Compared to their protein-coding counterparts, lncRNA genes are known to be less conserved due to lower evolutionary pressures.^23,24^ Accordingly, a prior analysis showed that there is only a 3.4% overlap between lncRNAs expressed in mouse and human among all tissues.^25^ Similarly, we found that only 0.88% of all lncRNAs were shared across both species. We found a smaller overlap than the prior study perhaps because we are focused only on lncRNAs expressed in the heart. None of the lncRNAs that were DE in TG compared to nTG mouse hearts were also DE in human HCM hearts compared to NF (Fig. 3B).

Among the 35 lncRNA genes that are DE in mice and possess a human ortholog, several of them have been shown by prior studies to be involved in cardiac physiology and pathophysiology. For instance, *ANRIL* acts as a transcriptional regular and epigenetic modifier in cardiovascular pathologies.^51-53^ *ANRIL* is antisense to *CDKN2B*, which codes for a cyclin-dependent kinase inhibitor, and thus can inhibit its expression. Accordingly, *ANRIL* KD has been shown to increase *CDKN2B* expression.^52,53^ Additionally, the human lncRNA *MIR181A1HG* has been shown to be a positive regulator of inflammation in atherosclerosis.^57,58^ However, no studies have linked *ANRIL* or *MIR181HG* specifically to HCM or cardiac hypertrophy. *ANRIL* and *MIR181HG* also have roles outside of cardiovascular biology such as cancer.^48-50^ Two of the other 35 lncRNAs, *E230013L22Rik* and *PARTICL*, have been studied before in the context of irradiation therapy and cancer,^44-47^ although they have no known cardiovascular function.

The GO analysis reveals that coding and non-coding gene sets related to non-coding RNA processing are significantly enriched. This suggests that non-coding RNA processing may be dysregulated in HCM pathophysiology. More specifically, the GSEA results for the human dataset show that genes involved in ncRNA processing are both upregulated and downregulated in HCM (Fig. 3C).

Many lncRNAs were falsely classified as non-coding by initial algorithms that arbitrarily set an open reading frame (ORF) threshold of <300 nucleotides in length to be considered non-coding. Recent work has identified that many ncRNAs contain sORFs that encode for micropeptides.^5,59,60^ We used the AlphaFold folding confidence metric pLDDT to assess for micropeptide coding potential since prior work has shown that sORFs from misannotated lncRNAs (i.e. lncRNAs that code for micropeptides) have high mean pLDDT values compared to random peptides and micropeptides for which there is high confidence that they are truly non-coding.^54^ We found this to be the case for three lncRNAs (*G730003C15Rik, 9830004L10Rik*, and *Gm45012*), indicating that they likely code for micropeptides. We found that two of those lncRNAs (*G730003C15Rik, 9830004L10Rik*) also had significantly lower mean PAE scores than controls. We observed alpha helical structures for the micropeptides coded *G730003C15Rik, 9830004L10Rik*, and *Gm4501*, which is a pattern also observed for misannotated lncRNAs in prior work.^54^ Lastly, positive PhyloCSF scores for two lncRNAs (*6330403L08Rik* and *2900072N19Rik*) indicate that they contain small (9-141 base pair) regions with phylogenetic patterns that are more likely in a coding model than a non-coding model.^22^ Indications of micropeptide coding potential for these lncRNAs provides a rationale for future work on HCM samples, which could identify sORFs from lncRNAs that are being actively translated into micropeptides.^4,5,60,61^

A limitation of our approach is that the human dataset comprised a smaller number of samples and a lower read depth (mean 57.32E6 (SD 7.03E6)) reads per sample)^14^ compared to the mouse dataset (77.87E6 (SD 9.45E6)). This likely limited the power to identify DE lncRNA genes across both species. Addressing this limitation could be prioritized for future studies on the contribution of lncRNAs to HCM pathophysiology. Additionally, a small amount of variance in the RNA-Seq data can be explained by genotype, as evidenced by PC1 and PC2 only capturing 8.95% of variance (Fig. 2B). This could indicate that a large amount of the variance is driven by unknown factors. Moreover, sex and exercise are covariates that we had to control for in order to capture as many DE lncRNAs as possible. Lastly, the control peptide sequences did not always have the expected pattern for pLDDT and PAE values. For instance, at range 1-10, the FNC positive control lncRNAs had a lower score than the TNC and random negative controls. This is the opposite of the expected behavior of the controls and thus the high score for *G730003C15Rik* at this range may not actually indicate coding potential.

In conclusion, we established a computational pipeline that identifies DE lncRNAs in a murine disease model, predicts human orthologs, merges analyses between mouse and human RNA-Seq data sets, and quantifies micropeptide coding potential. Using this pipeline, we identified lncRNAs that were both downregulated and upregulated in a mouse model of HCM, several of which had human orthologs and can be prioritized for future experimental studies to probe their role in HCM pathophysiology.

## DATA AVAILABILITY

Mouse and human RNA-Seq data is from GSE297707 and GSE130036, respectively.

## SUPPLEMENTAL MATERIAL

Supplemental Figures S1-3, Table T1, and Code C1 can be found at: https://github.com/gbranscom/lncrna_hcm/tree/main (DOI: https://doi.org/10.6084/m9.figshare.29604356.v1).

## ACKNOWLEDGMENTS

We would like to acknowledge Taehyong Kim, Jingjing Li, Yifan Yang, Marcus Wagner, Christopher McAllister, Hyung Chul Kim, Emmitt Jolly, Dmitry Mylarshchikov, and the Penn Medicine Academic Computing Services staff for their technical assistance.

## AUTHOR CONTRIBUTIONS

G.A.B. performed experiments, analyzed the data, interpreted the results of experiments, prepared figures, and drafted the manuscript. S.M.D. conceived and designed the study, along with editing, revising, and approving the final version of the manuscript. J.J.H. conceived the initial mouse study and performed the RNA isolation and sequencing. M.M. provided input into the bioinformatics approaches. J.M.Y., M.M., and J.J.H. provided technical support and critical review of the manuscript.

