## Supplemental Figures S1-3, Table T1, and Code C1 for "Investigating the role of long non-coding RNA in hypertrophic cardiomyopathy": cufflinks.sh

```
# transcript essembly with cufflinks

# 1. cufflinks transcript essembly

$reference_dir="/path/to/dir/with/reference/files"
$in_dir="/path/to/input/bam/files"
$out_dir="/path/to/out/dir"

for folder_path in "$in_dir"/*/; do
	cd "$folder_path"

	unique_id=$(basename $(pwd))
    cufflinks \
    -p 60 \
    --library-type fr-firststrand \
    -g ${reference_dir}/gencode.vM10.primary_assembly.annotation.gtf \
    -o $out_dir \
    ${unique_id}.bam

    cd ..
done

# 2. merge results into single GTF file with cuffmerge

$MANIFEST_FILE="/path/to/manifest_file.txt"
cd $out_dir

touch "$MANIFEST_FILE" # create manifest_file.txt with list of individual GTF files
for file in "$out_dir"/*.gtf; do
    echo "$file" >> "$MANIFEST_FILE" 
done

cuffmerge \
-p 60 \
-s ${reference_dir}/GRCm38.primary_assembly.genome.fa \
-g ${reference_dir}/gencode.vM10.primary_assembly.annotation.gtf \
./manifest.txt

# this output file is provided (lncrna.csv)
```
