## Supplemental Figures S1-3, Table T1, and Code C1 for "Investigating the role of long non-coding RNA in hypertrophic cardiomyopathy": cutadapt.sh

```
# adapter trimming using cutadapt for all fastq files

$in_dir="/path/to/input/dir" # fastq files
$out_dir="/path/to/output/dir"

for folder_path in "$input_dir"/*/; do
    cd "$folder_path"
    unique_id=$(basename $(pwd))
    mkdir $out_dir/$unique_id
    cutadapt -j=40 \
        -m 1 \
        -a AGATCGGAAGAGCACACGTCTGAACTCCAGTCA \
        -A AGATCGGAAGAGCGTCGTGTAGGGAAAGAGTGT \
        -o $out_dir/$unique_id/{$unique_id}_R1_001_trimmed.fastq.gz \
        -p $out_dir/$unique_id/{$unique_id}_R2_001_trimmed.fastq.gz \
        ./*_R1_001.fastq.gz \
        ./*_R2_001.fastq.gz
    cd ..
done
```
