## Supplemental Figures S1-3, Table T1, and Code C1 for "Investigating the role of long non-coding RNA in hypertrophic cardiomyopathy": micropeptide.sh

```
# predicts micropeptides for lncRNAs

# 1. Predict sORFs from gene sequences using MiPepid
# download source code here: https://github.com/MindAI/MiPepid
# prequisites: combined_exons_seqs.fa (FASTA file where each record contains a different contiguos exon sequence)

mipepid_dir="$path/to/mipepid/dir"
cd $mipepid_dir
python ./src/mipepid.py $mipepid_dir/combined_exon_seqs.fa $mipepid_dir/outfile.csv

# 2. Translate sORF nucletoide sequences into peptide sequences with EMBOSS transeq
# prerequisites: create separate FASTA file for each sORF

$in_dir="$path/to/nucleotide/seq/fasta/files/dir"
for file in "$in_dir"/*; do
    filename=$(basename "$file")
    transeq -trim $file "${out_dir}/${filename}"
done

# 3. Run AlphaFold2 on each sORF's FASTA file
# download AlphaFold2 locally or run on BATCH: https://colab.research.google.com/github/sokrypton/ColabFold/blob/main/batch/AlphaFold2_batch.ipynb
```
