## Supplemental Figures S1-3, Table T1, and Code C1 for "Investigating the role of long non-coding RNA in hypertrophic cardiomyopathy": rsem.sh

```
# generation of count matrices with RSEM

# 1. generate reference

$reference_dir="/path/to/dir/with/reference/files"
$star="/path/to/star/installation"
$rsem_reference_dir="/path/to/rsem/ref/dir"

rsem-prepare-reference \
    --star \
    --star-path $star \
    -p 60 \
    --gtf /path/to/cuffmerge_output.gtf \
    ${reference_dir}/GRCm38.primary_assembly.genome.fa \
    $rsem_reference_dir

# 2. generate count matrices

$in_dir="/path/to/input/dir" # fastq files

for folder_path in "$input_dir"/*/; do
	cd "$folder_path"

	unique_id=$(basename $(pwd))
    rsem-calculate-expression \
        --paired-end \
        --strandedness reverse \
        -p 20 \
        --star \
        --star-path $star \
        --star-gzipped-read-file \
        --sort-bam-by-coordinate \
        --output-genome-bam \
        --sort-bam-memory-per-thread 40G \
        ${in_dir}/*_R1_* \
        ${in_dir}/*_R2_* \
        $rsem_reference_dir \
        $unique_id

    cd ..
done
```
