## Supplemental Figures S1-3, Table T1, and Code C1 for "Investigating the role of long non-coding RNA in hypertrophic cardiomyopathy": star.sh

```
# genome alignment with STAR

# 1. generate indices

$genome_dir="/path/to/genome/dir"
$reference_dir="/path/to/dir/with/reference/files"

$HOME/bin/STAR \
--runThreadN 40 \
--runMode genomeGenerate \
--genomeDir $genome_dir \
--genomeFastaFiles ${reference_dir}/GRCm38.primary_assembly.genome.fa \
--sjdbGTFfile ${reference_dir}/gencode.vM10.primary_assembly.annotation.gtf \
--sjdbOverhang 150

# 2. align reads

for folder_path in "$trimmed_fastq_dir"/*/; do
	cd "$folder_path"
	unique_id=$(basename $(pwd))
    STAR \
    --runMode alignReads \
    --runThreadN 40 \
    --genomeDir $genome_dir \
    --readFilesIn ${unique_id}_R1_001.fastq.gz ${unique_id}_R2_001.fastq.gz \
    --outSAMtype BAM SortedByCoordinate \
    --readFilesCommand gunzip -c \
    --outFileNamePrefix "path/to/out/dir/${unique_id}"
    cd ..
done
```
