## Supplemental Figures S1-3, Table T1, and Code C1 for "Investigating the role of long non-coding RNA in hypertrophic cardiomyopathy": lncrna_deseq.Rmd

````
# DESeq for mouse lncRNAs
  
# Description/Goals

DESeq analaysis for lncRNA candidates identified from the GENCODE lncRNA GTF and any extra novel lncRNAs from FEELnc.

We use the gene-level abundance estimates from RSEM and `tximport` to obtain the count matrix.

# Prerequisites

* lncrna.csv: lncRNA GTF file (Cuffmerge output) converted to CSV

# Data prep

## General imports

```{r}
if(!require('pacman')) {install.packages('pacman')}
pacman::p_load(tidyverse, ggplot2, dplyr, ggrepel, gridExtra, plotly, sunburstR, d3r, pheatmap, devtools, xlsx, ggVennDiagram)
```

## Imports for DESeq2

```{r}
if (!requireNamespace("BiocManager", quietly = TRUE)) {
  install.packages("BiocManager")
}
install_if_missing <- function(pkg) {
  if (!requireNamespace(pkg, quietly =TRUE)) {
    BiocManager::install(pkg)
  }
}
bioc.packages <- c("tximport", "tximportData", "DESeq2", "apeglm", "DEGreport",
                   "vsn", "EnhancedVolcano", "org.Mm.eg.db")
lapply(bioc.packages, install_if_missing)
```

## lncRNA input data

```{r}
lncrna_df <- read.csv("./data/lncrna.csv")
```

## tximport

[tximport docs](https://bioconductor.org/packages/devel/bioc/vignettes/tximport/inst/doc/tximport.html#RSEM)

```{r, echo=TRUE, results=FALSE}
# locate the directory for the `tximportData` files
# contents of dir: subdirs with name of each sample, inside each subdir is RSEM output file (samplename.genes.results )
dir <- system.file("extdata", package = "tximportData")

# read in metadata for each sample
treatments.df <- read.csv("./data/metadata.csv") # columns: sample, genotype, exercise, sex

# generate gene-level count matrix
gene.files <- file.path(dir, treatments.df$sample, paste0(samples$sample, ".genes.results"))
names(gene.files) <- sapply(gene.files, treatments.df$sample)
txi.genes.rsem <- tximport(gene.files, type = "rsem", txIn = FALSE, txOut = FALSE)
txi.genes.rsem.df <- as.data.frame(txi.genes.rsem$counts)
```
            
# Conduct DESEq2

```{r}
# individual variables
dds.genes.individual <- DESeqDataSetFromTximport(txi = txi.genes.rsem,
                                      colData = col.data,
                                      design = ~ genotype + exercise + sex)
dds.genes.individual <- DESeq(dds.genes.individual)
dds.genes.individual
plotDispEsts(dds.genes.individual)

# disaggregated analyses

# combined genotype_exercise
dds.genes.combined.1 <- DESeqDataSetFromTximport(txi = txi.genes.rsem,
                                      colData = col.data,
                                      design = ~ genotype_exercise + sex)
dds.genes.combined.1 <- DESeq(dds.genes.combined.1)
dds.genes.combined.1
plotDispEsts(dds.genes.combined.1)

# combined genotype_sex + exercise
dds.genes.combined.2 <- DESeqDataSetFromTximport(txi = txi.genes.rsem,
                                      colData = col.data,
                                      design = ~ genotype_sex + exercise)
dds.genes.combined.2 <- DESeq(dds.genes.combined.2)
dds.genes.combined.2
plotDispEsts(dds.genes.combined.2)

# define p-value cutoff
pcut = 0.05

# TG vs nTG
res_TG_vs_nTG <- results(dds.genes.individual, contrast=c("genotype","TG","nTG"), lfcThreshold=fccut, altHypothesis="greaterAbs", pAdjustMethod="BH", independentFiltering = TRUE, alpha=pcut, tidy=TRUE)
res_ordered_TG_vs_nTG <- res_TG_vs_nTG[order(res_TG_vs_nTG$pvalue,
                                             -abs(res_TG_vs_nTG$log2FoldChange)),]
res_ordered_TG_vs_nTG

# M vs F
res_M_vs_F <- results(dds.genes.individual, contrast=c("sex","m","f"), lfcThreshold=fccut, altHypothesis="greaterAbs", pAdjustMethod="BH", independentFiltering = TRUE, alpha=pcut, tidy=TRUE)
res_M_vs_F

# HIIT vs Sed
res_HIIT_vs_Sed <- results(dds.genes.individual, contrast=c("exercise","HIIT","Sed"), lfcThreshold=fccut, altHypothesis="greaterAbs", pAdjustMethod="BH", independentFiltering = TRUE, alpha=pcut, tidy=TRUE)
res_HIIT_vs_Sed

# TG_Sed vs nTG_Sed
res_TG_Sed_vs_nTG_Sed <- results(dds.genes.combined.1, contrast=c("genotype_exercise","TG_Sed","nTG_Sed"), lfcThreshold=fccut, altHypothesis="greaterAbs", pAdjustMethod="BH", independentFiltering = TRUE, alpha=pcut, tidy=TRUE)
res_ordered_TG_Sed_vs_nTG_Sed <- res_TG_Sed_vs_nTG_Sed[order(res_TG_Sed_vs_nTG_Sed$pvalue,
                                                             -abs(res_TG_Sed_vs_nTG_Sed$log2FoldChange)),]
rownames(res_ordered_TG_Sed_vs_nTG_Sed) <- NULL
res_ordered_TG_Sed_vs_nTG_Sed

# TG_HIIT vs nTG_HIIT
res_TG_HIIT_vs_nTG_HIIT <- results(dds.genes.combined.1, contrast=c("genotype_exercise","TG_HIIT","nTG_HIIT"), lfcThreshold=fccut, altHypothesis="greaterAbs", pAdjustMethod="BH", independentFiltering = TRUE, alpha=pcut, tidy=TRUE)
res_ordered_TG_HIIT_vs_nTG_HIIT <- res_TG_HIIT_vs_nTG_HIIT[order(res_TG_HIIT_vs_nTG_HIIT$pvalue,
                                                                 -abs(res_TG_HIIT_vs_nTG_HIIT$log2FoldChange)),]
rownames(res_ordered_TG_HIIT_vs_nTG_HIIT) <- NULL
res_ordered_TG_HIIT_vs_nTG_HIIT

# TG_m_vs_nTG_m
TG_m_vs_nTG_m <- results(dds.genes.combined.2, contrast=c("genotype_sex","TG_m","nTG_m"), lfcThreshold=fccut, altHypothesis="greaterAbs", pAdjustMethod="BH", independentFiltering = TRUE, alpha=pcut, tidy=TRUE)
res_ordered_TG_m_vs_nTG_m <- res_TG_m_vs_nTG_m[order(res_TG_m_vs_nTG_m$pvalue,
                                                                 -abs(res_TG_m_vs_nTG_m$log2FoldChange)),]
rownames(res_ordered_TG_m_vs_nTG_m) <- NULL
res_ordered_TG_m_vs_nTG_m

# TG_f_vs_nTG_f
TG_f_vs_nTG_f <- results(dds.genes.combined.2, contrast=c("genotype_sex","TG_f","nTG_f"), lfcThreshold=fccut, altHypothesis="greaterAbs", pAdjustMethod="BH", independentFiltering = TRUE, alpha=pcut, tidy=TRUE)
res_ordered_TG_f_vs_nTG_f <- res_TG_f_vs_nTG_f[order(res_TG_f_vs_nTG_f$pvalue,
                                                                 -abs(res_TG_f_vs_nTG_f$log2FoldChange)),]
rownames(res_ordered_TG_f_vs_nTG_f) <- NULL
res_ordered_TG_f_vs_nTG_f

# variance stabilizing stabilization (vsd)
vsd.genes.individual <- varianceStabilizingTransformation(dds.genes.individual, blind=FALSE)
vsd.genes.combined.1 <- varianceStabilizingTransformation(dds.genes.combined.1, blind=FALSE)
vsd.genes.combined.2 <- varianceStabilizingTransformation(dds.genes.combined.2, blind=FALSE)
meanSdPlot(assay(vsd.genes.individual))
meanSdPlot(assay(vsd.genes.combined.1))
meanSdPlot(assay(vsd.genes.combined.2))

# add gene names and info from lncrna_df
lncrna_df <- read.csv("./data/lncrna.csv") # provided
add.gene.names.helper <- function(df.filtered, df.name) {
  df.merged <- merge(df.filtered, lncrna_df, by="gene_name")
  other.cols <- setdiff(colnames(df.merged), c("gene_name", "gene_id"))
  df.merged <- df.merged %>% dplyr::select('gene_name', 'gene_id', all_of(other.cols))
  return(df.merged)
}

add.gene.names <- function() {
  type <- tolower(type)
  
  dfs <- list(
    list(res_TG_vs_nTG, "res_TG_vs_nTG_filtered"),
    list(res_TG_Sed_vs_nTG_Sed, "res_TG_Sed_vs_nTG_Sed_filtered"),
    list(res_TG_HIIT_vs_nTG_HIIT, "res_TG_HIIT_vs_nTG_HIIT_filtered"),
    list(res_TG_m_vs_nTG_m, "res_TG_m_vs_nTG_m_filtered"),
    list(res_TG_f_vs_nTG_f, "res_TG_f_vs_nTG_f_filtered")
    )
  
  for (i in 1:length(dfs)) {
    df <- dfs[[i]][[1]]
    name <- dfs[[i]][[2]]
    
    df.filtered <- df %>%
      filter(padj <= pcut) %>%
         arrange(padj, desc(abs(log2FoldChange)))
    assign(name, create.merged.helper(df.filtered, name), envir = .GlobalEnv)
  }
}
```

# Data visualization

## Preliminary data exploration

```{r}
# heatmap of count matrix
select <- order(rowMeans(counts(dds.genes.individual,normalized=TRUE)),
                decreasing=TRUE)[1:40] # only include the top 40 genes
df <- as.data.frame(colData(dds.genes.individual)[,c("genotype", "exercise", "sex")])
pheatmap(assay(vsd.genes.individual)[select,], cluster_rows=FALSE, show_rownames=FALSE,
         cluster_cols=FALSE, annotation_col=df, fontsize_col=6, scale="row",
         main="heatmap of count matrix:\n top 40 counts for each sample - variance stabilizing stabilization")

# PCA
gene_var <- rowVars(assay(vsd.genes.individual))
top_var_genes <- order(gene_var, decreasing = TRUE)[1:500]
tr <- t(assay(results)[top_var_genes, ])
pca <- prcomp(tr)
names <- names(pca)
loadings <- pca$rotation
values <- pca$x
pve <- summary(pca)$importance[2, ]
pc1_var <- round(pve[1] * 100, 2)
pc2_var <- round(pve[2] * 100, 2)

# colored by genotype
pca.genotype <- as.data.frame(pca$x) %>% 
  ggplot(aes(x=PC1, y=PC2)) +
  geom_point(aes(color = results$genotype)) +
  geom_vline(xintercept = 0) +
  geom_hline(yintercept = 0) +
  labs(
    x = paste0("PC1 (", pc1_var, "% variance)"),
    y = paste0("PC2 (", pc2_var, "% variance)"),
    color = "genotype"
    ) +
  theme_minimal() +
  scale_color_manual(values = c(
    "nTG" = "#68cdc4",
    "TG" = "#7a66a8"
  )) +
  ggtitle(sprintf("PC1 vs. PC2 for %s - color by genotype", name))
print(pca.genotype)

# colored by exercise group
pca.exercise <- as.data.frame(pca$x) %>% 
    ggplot(aes(x=PC1, y=PC2)) +
    geom_point(aes(color = results$exercise)) +
    geom_vline(xintercept = 0) +
    geom_hline(yintercept = 0) +
    labs(
      x = paste0("PC1 (", pc1_var, "% variance)"),
      y = paste0("PC2 (", pc2_var, "% variance)"),
      color = "exercise"
      ) +
    theme_minimal() +
    ggtitle(sprintf("PC1 vs. PC2 for %s - color by exercise group", name))
print(pca.exercise)
  
# colored by sex
pca.sex <- as.data.frame(pca$x) %>% 
ggplot(aes(x=PC1, y=PC2)) +
  geom_point(aes(color = results$sex)) +
  geom_vline(xintercept = 0) +
  geom_hline(yintercept = 0) +
  labs(
    x = paste0("PC1 (", pc1_var, "% variance)"),
    y = paste0("PC2 (", pc2_var, "% variance)"),
    color = "sex"
    ) +
  theme_minimal() +
  scale_color_manual(values = c(
    "m" = "lightblue",
    "f" = "pink"
  )) +
  ggtitle(sprintf("PC1 vs. PC2 for %s - color by sex", name))
print(pca.sex)
```

## Volcano plots

```{r}
# TG vs nTG
EnhancedVolcano(res_TG_vs_nTG,
  lab = res_TG_vs_nTG$gene_name,
  x = 'log2FoldChange',
  y = 'padj',
  title = 'DE lncRNA genes: TG vs nTG',
  pCutoff = pcut,
  FCcutoff = 0,
  pointSize = 3.0,
  labSize = 6.0)

# M vs F
EnhancedVolcano(res_M_vs_F,
  lab = res_M_vs_F$gene_name,
  x = 'log2FoldChange',
  y = 'padj',
  title = 'DE lncRNA genes: M vs F',
  pCutoff = pcut,
  FCcutoff = 0,
  pointSize = 3.0,
  labSize = 6.0)

# HIIT vs Sed
EnhancedVolcano(res_HIIT_vs_Sed,
  lab = res_HIIT_vs_Sed$gene_name,
  x = 'log2FoldChange',
  y = 'padj',
  title = 'HIIT vs Sed',
  pCutoff = pcut,
  FCcutoff = fccut,
  pointSize = 3.0,
  labSize = 6.0)

# TG_Sed vs nTG_Sed
EnhancedVolcano(res_TG_Sed_vs_nTG_Sed,
  lab = res_TG_Sed_vs_nTG_Sed$gene_name,
  x = 'log2FoldChange',
  y = 'padj',
  title = 'TG_Sed vs nTG_Sed',
  pCutoff = pcut,
  FCcutoff = fccut,
  pointSize = 3.0,
  labSize = 6.0)

# TG_HIIT vs nTG_HIIT
EnhancedVolcano(res_TG_HIIT_vs_nTG_HIIT,
  lab = res_TG_HIIT_vs_nTG_HIIT$gene_name,
  x = 'log2FoldChange',
  y = 'padj',
  title = 'TG_HIIT vs nTG_HIIT',
  pCutoff = pcut,
  FCcutoff = fccut,
  pointSize = 3.0,
  labSize = 6.0
  )

# TG_m vs nTG_m
EnhancedVolcano(res_TG_m_vs_nTG_m,
  lab = res_TG_m_vs_nTG_m$gene_name,
  x = 'log2FoldChange',
  y = 'padj',
  title = 'TG_m vs nTG_m - q-value',
  pCutoff = pcut,
  FCcutoff = 0,
  pointSize = 3.0,
  labSize = 6.0
)

# TG_f vs nTG_f
EnhancedVolcano(res_TG_f_vs_nTG_f,
  lab = res_TG_f_vs_nTG_f$gene_name,
  x = 'log2FoldChange',
  y = 'padj',
  title = 'TG_f vs nTG_f - q-value (BH-corrected)',
  pCutoff = pcut,
  FCcutoff = 0,
  pointSize = 3.0,
  labSize = 6.0
  )
```
````
