## Supplemental Figures S1-3, Table T1, and Code C1 for "Investigating the role of long non-coding RNA in hypertrophic cardiomyopathy": README.md

# lncrna_hcm

Supplementary code C1 for the manuscript: "Investigating the role of long non-coding RNA in hypertrophic cardiomyopathy"

> Paper citation TBD

Supplementary Figures S1-3 and Supplementary Table T1 are also found here.

## Overview

We implemented a complete RNA-seq analysis pipeline including:

- **Adapter trimming** using `cutadapt`
- **Read alignment** with `STAR`
- **Transcript assembly and quantification** with `Cufflinks` and `RSEM`
- **Differential expression (DE) analysis** using `DESeq2` in R
- **Prediction of lncRNA-encoded micropeptides** using `MiPepid` and `AlphaFold2`

This code is supplementary material that can be used to reproduce our findings.

## Repository Structure
```text
├── reference_files.txt     # List of reference genome files
├── environment.yml         # Conda environment definition
├── metadata.csv            # Sample metadata on mouse samples
├── lncrna.csv              # GTF output of Cuffmerge (all lncRNAs captured by RNA-Seq among all 64 mice)
├── de_seq.Rmd              # R markdown document for DESeq
└── README.md               # This file
├── scripts/                # Bash scripts
├── supplementary_material/ # Supplementary Figures S1-3 and Supplementary Table T1
```

## Getting Started

### 1. Clone this repository

```bash
git clone https://github.com/gbranscom/lncrna_hcm.git
cd lncrna_hcm
```

### 2. Set up the conda environment

```bash
conda env create -f environment.yml
conda activate lncrna_hcm
```

### 3. Download the reference files

List of reference files can be found in `reference_files.txt`.

### 4. Run the pipeline for mouse RNA-Seq

```bash
cutadapt.sh # 1. Adapter trimming
star.sh # 2. Genome alignment
cufflinks.sh # 3. Transcript assembly
rsem.sh # 4. Count matrix generation
lncrna_deseq.R # 5. Differential expression analysis
micropeptide.sh # 6. Micropeptide prediction
```

Information on the human DE analysis can be found in

> Liu X, Ma Y, Yin K, Li W, Chen W, Zhang Y, et al. Long non-coding and coding RNA profiling using strand-specific RNA-seq in human hypertrophic cardiomyopathy. Scientific Data. 2019 Jun 13;6(1). doi:10.1038/s41597-019-0094-6

## Questions?

Please reach out to me with any issues: graham.branscom 'at' gmail.com
