## Supplemental Figures S1-3, Table T1, and Code C1 for "Investigating the role of long non-coding RNA in hypertrophic cardiomyopathy": supp_figure_legends.docx

Supplemental Figure S1. Volcano plots from the TG vs nTG analysis stratified by sex and exercise group

A. DE lncRNAs in *TNNT2* ∆160 transgenic (TG) vs non-transgenic (nTG) males only. *y*-axis denotes *p*-values adjusted with the Benjamini-Hochberg procedure. The horizontal line above the x-axis indicates a *p*-adjusted cutoff of 0.05.
B. Same as panel A but for females only.
C. Same as panel A but for HIIT mice only.
D. Same as panel A but for sedentary mice only

TG: transgenic, nTG: nontransgenic, M: male, F: female, HIIT: high intensity interval training, Sed: sedentary

Supplemental Figure S2. Number of sORFs per lncRNA and AlphaFold PAE plot

A. Number of small open reading frames (sORFs) for each of the 13 DE (*p*-adj < 0.05) mouse lncRNAs which have a human ortholog. The statistical significance of each number is compared to the mean number of sORFs among all mouse lncRNA transcripts (n=155,417). Error bars depict standard deviation. *p*-values were determined via permutation test.
B. Measures of folding confidence for micropeptides coded by the 13 DE (*p*-adj < 0.05) mouse lncRNAs with human orthologs, stratified by micropeptide length. Only lncRNA genes with significant results are shown. Mean per-residue predicted aligned error (PAE) scores are for micropeptides falling into buckets of length 10 amino acids long from 1 to 80. Lower PAE scores indicate lower folding confidence and higher micropeptide coding potential. There are n=70 micropetides for *G730003C15Rik* and n=233 for *9830004L10Rik*. There are 120 micropeptides each for false non-coding (FNC, i.e. true coding), random, and true non-coding (TNC), 15 for each length range. Error bars depict standard deviation. *p*-values were determined with independent two sample t-tests. *p*-values are denoted by asterisks sitting above and having the color of the two groups that they are comparing within a given length range subset. For simplicity, *p*-values are only shown when any of the three lncRNAs have a greater pLDDT than random or TNC micropeptides.

* *p* < 0.05, ** p < 0.01, *** p < 0.001, **** p < 0.0001

sORF: small open reading frame, PAE: predicted alignment error, FNC: false non-coding (i.e. true coding), TNC: true non-coding

Supplemental Figure S3. Additional PhyloCSF result

PhyloCSF tracks in the UCSC Genome Browser for the mouse lncRNA *2900072N19Rik* in genome build GRCm38/mm10. The topmost track is the PhyloCSF novel track, indicating novel coding regions (green is on plus strand, red is on minus strand). Below that is the Smoothed PhyloCSF track for each of the six possible reading frames, which depicts the scores given by PhyloCSF’s hidden Markov model. Next is the PhyloCSF regions track for all reading frames, indicating the Smoothed track’s predicted coding regions. Next is the PhyloCSF Power track, which indicates a score from 0 to 1 indicating statistical confidence, with a higher score indicating higher confidence. Next are the genes located within this region. *6330403L08Rik* is marked with a red arrow. Lastly, there is the Conservation track of 100 vertebrates, including mouse, for reference.
