## Supplemental Figures S1-3, Table T1, and Code C1 for "Investigating the role of long non-coding RNA in hypertrophic cardiomyopathy": supp_table_1.pdf

**Supplemental Table 1:** lncRNA genes that are DE in mouse.

| Mouse (GRCm38/mm10) | | | | | Human (GRCh38/hg19) | | | |
|---|---|---|---|---|---|---|---|---|
| **lncRNA gene name (Ensembl ID)** | **comparison source** | **$log_2FC$** | ***p*-adj** | **known tissue expression** | **lncRNA ortholog gene name (Ensembl ID)** | **FC info** | **known tissue expression** | **known function** |
| Gm43672 (ENSMUSG00000106019) | TG vs nTG | -0.557 | 7.62E-11 | Heart, brain, muscle, trachea, skin, spleen<br><br>https://rnacentral.org/rna/URS0002A33156/10090 | ENSG00000308496 | | | |
| | TG_f vs nTG_f | -0.630 | 1.04E-06 | | | | | |
| | TG_HIIT vs nTG_HIIT | -0.583 | 3.17E-06 | | | | | |
| | TG_Sed vs nTG_Sed | -0.531 | 6.55E-04 | | | | | |
| | TG_m vs nTG_m | -0.486 | 1.99E-03 | | | | | |
| Gm9905 (ENSMUSG00000053358) | TG vs nTG | -0.594 | 3.10E-08 | | | | | |
| | TG_HIIT vs nTG_HIIT | -0.689 | 3.17E-06 | | | | | |
| | TG_m vs nTG_m | -0.587 | 1.74E-03 | | | | | |
| | TG_f vs nTG_f | -0.602 | 2.28E-03 | | | | | |
| Gm20632 (ENSMUSG00000093577) | TG_HIIT vs nTG_HIIT | -0.546 | 1.80E-03 | All tissues have low expression<br><br>https://rnacentral.org/rna/URS0002A237A7/10090 | ENSG00000290792 | | | |
| Gm42918 (ENSMUSG00000106209) | TG_m vs nTG_m | -0.951 | 9.07E-04 | | | | | |
| | TG vs nTG | -0.680 | 1.20E-03 | | | | | |
| | TG_HIIT vs nTG_HIIT | -0.810 | 4.42E-03 | | | | | |
| Banf2os (ENSMUSG00000086384) | TG vs nTG | -0.965 | 3.11E-08 | All tissues have low expression<br><br>https://rnacentral.org/rna/URS0002A19B47/10090 | | | | |
| | TG_f vs nTG_f | -1.086 | 1.38E-04 | | | | | |
| | TG_Sed vs nTG_Sed | -1.047 | 6.55E-04 | | | | | |
| | TG_HIIT vs nTG_HIIT | -0.883 | 1.80E-03 | | | | | |
| | TG_m vs nTG_m | -0.842 | 1.01E-02 | | | | | |
| Gm13375 (ENSMUSG00000075514) | TG_HIIT vs nTG_HIIT | -0.488 | 4.46E-03 | Brain, skin, bone, thymus, lung, colon, spleen<br><br>https://rnacentral.org/rna/URS0002A2504A/10090 | | | | |
| RP23-166L22.2 (ENSMUSG00000110340) | TG vs nTG | -1.361 | 1.64E-06 | All tissues have low expression | | | | |
| | TG_f vs nTG_f | -1.740 | 5.94E-03 | | | | | |

| | | | | | | | | |
|---|---|---|---|---|---|---|---|---|
| | TG_m vs nTG_m | -1.159 | 1.01E-02 | | | | | |
| G730003C15Rik (ENSMUSG00000097573) | TG_HIIT vs nTG_HIIT | -0.684 | 5.47E-03 | https://rnacentral.org/rna/URS0002A4F318/10090 | ENSG00000287133 | | | |
| | TG vs nTG | -0.536 | 8.45E-03 | | | | | |
| E230013L22Rik (ENSMUSG00000096957) | TG vs nTG | -0.476 | 9.14E-05 | Heart<br><br>https://rnacentral.org/rna/URS0002A1C2B5/10090 | | | | Most upregulated lncRNA after X-ray irradiation in mouse lung tissue, modulates the mRNA BNIP1[44] |
| | TG_HIIT vs nTG_HIIT | -0.539 | 1.80E-03 | | | | | |
| | TG_f vs nTG_f | -0.525 | 2.13E-02 | | | | | |
| 2900072N19Rik (ENSMUSG00000087018) | TG vs nTG | -0.760 | 8.45E-03 | Brain<br><br>https://rnacentral.org/rna/URS0002A5782B/10090 | LINC02710 (ENSG00000255269) | | Testis<br><br>https://rnacentral.org/rna/URS0000EF618F/9606 | |
| C530005A16Rik (ENSMUSG00000085408) | TG vs nTG | -0.414 | 9.85E-03 | Brain<br><br>https://rnacentral.org/rna/URS0002A5A50C/10090 | | | | |
| Prr33 (ENSMUSG00000043795) | TG_HIIT vs nTG_HIIT | -0.519 | 1.80E-03 | | | | | Annotated in GENCODE M10 as long intergenic non-coding RNA, but traditionally annotated as protein-coding. |
| | TG_m vs nTG_m | -0.499 | 1.01E-02 | | | | | |
| | TG vs nTG | -0.373 | 2.68E-02 | | | | | |
| Gm37752 (ENSMUSG00000103385) | TG vs nTG | 2.438 | 1.83E-02 | | | | | |
| Gm45012 (ENSMUSG00000109052) | TG vs nTG | -0.422 | 9.49E-04 | Heart<br><br>https://rnacentral.org/rna/URS0002A2D746/10090 | ENSG00000300258 | | | |
| | TG_HIIT vs nTG_HIIT | -0.415 | 4.00E-02 | | | | | |
| PARTICL (ENSG00000286532) | TG_HIIT vs nTG_HIIT | -0.468 | 2.45E-02 | Heart, arteries, prostate, esophagus, brain, fallopian tubes, cervix, testis, colon, thyroid, bladder, skin<br><br>https://rnacentral.org/rna/URS0002A47137/9606 | PARTICL (ENSG00000286532) | | | Upregulated following low-dose irradiation. Alters H3K27me3 distribution and interacts with DNA methyltransferase 1.[45,46] Forms triple helices clustered at tumor suppressor genes.[47] |
| Mir223hg (ENSMUSG00000078122) | TG_m vs nTG_m | 0.685 | 1.99E-03 | Heart, lung, Spleen, liver, skin, mammary gland, bone, uterus, | Mir223hg (ENSG00000274536) | | Appendix, tonsil, lung, bladder, bone, leukocyte, lymph node, | Positive regulator of inflammation in atherosclerosis.[57,58] Differentially expressed in lung adenocarcinoma and ovarian cancer.[48,49] |
| | TG_HIIT vs nTG_HIIT | 0.520 | 4.72E-02 | | | | | |

| | | | | | | | | |
|---|---|---|---|---|---|---|---|---|
| | | | | bladder, adrenal gland,<br><br>https://rnacentral.org/rna/URS0002A33BA1/10090 | | | small intestine, spleen<br><br>https://rnacentral.org/rna/URS0002A35E6A/9606 | |
| Gm28703 (ENSMUSG00000099512) | TG_m vs nTG_m<br>TG vs nTG | -0.616<br>-0.470 | 2.11E-02<br>2.89E-02 | All tissues have low expression<br><br>https://rnacentral.org/rna/URS0002A2C79C/10090 | | | | |
| Gm10435 (ENSMUSG00000072902) | TG vs nTG<br>TG_m vs nTG_m<br>TG_HIIT vs nTG_HIIT | -0.389<br>-0.428<br>-0.387 | 1.93E-03<br>3.38E-02<br>4.40E-02 | Heart, lung,<br><br>https://rnacentral.org/rna/URS000078648E/10090 | ENSG00000287315 | | | |
| 9830004L10Rik (ENSMUSG00000099552) | TG vs nTG | -0.660 | 2.68E-02 | | ENSG00000300443 | | | |
| Gm7628 (ENSMUSG00000025644) | TG vs nTG | -0.731 | 2.68E-02 | All tissues have low expression<br><br>https://rnacentral.org/rna/URS0000811860/10090 | | | | |
| Gm17586 (A_J_v1:CM003964.1) | TG vs nTG<br>TG_HIIT vs nTG_HIIT<br>TG_m vs nTG_m | 0.759<br>0.691<br>0.758 | 2.69E-04<br>4.40E-02<br>4.79E-02 | | | | | |
| ANRIL (ENSG00000240498) | TG vs nTG | 0.649 | 3.10E-02 | Small intestine, colon, brain<br><br>https://rnacentral.org/rna/URS00026A21AD/9606 | ANRIL (ENSG00000264545) | | | Transcriptional regulator and epigenetic modifier in myocardial infarction and coronary artery disease.[51] Antisense to CDKN2B. ANRIL KD has been shown to increase CDKN2B expression.[52,53] Biomarker of different caners.[50] |
| 6330403L08Rik (ENSMUSG00000075585) | TG_HIIT vs nTG_HIIT<br>TG_HIIT vs nTG_HIIT<br>TG_HIIT vs nTG_HIIT | -0.387<br>-0.387<br>-0.387 | 3.14E-02<br>3.14E-02<br>3.14E-02 | Heart, brain, skeletal muscle, colon, small intestine, trachea<br><br>https://rnacentral.org/rna/URS0000775964/10090 | PDGFA-DT (ENSG00000223855) | HCM_m vs ctrl_m: log2FC 0.327, p-adj 2.62E-01<br><br>HCM_f vs ctrl_f: log2FC -0.499, p-adj 6.38E-01<br><br>HCM vs ctrl: log2FC 0.194, p-adj 5.17E-01 | Heart<br><br>https://rnacentral.org/rna/URS000314D59/9606 | |

| | | | | | | | |
|---|---|---|---|---|---|---|---|
| Gm45051 (ENSMUSG00000108435) | TG_HIIT vs nTG_HIIT | -0.582 | 3.14E-02 | Kidney<br><br>https://rnacentral.org/rna/URS00009C60DF/10090 | | | |
| Gm10115 (ENSMUSG00000062319) | TG_HIIT vs nTG_HIIT<br>TG vs nTG | 0.540<br>0.443 | 3.14E-02<br>3.48E-02 | Heart, brain<br><br>https://rnacentral.org/rna/URS0000781B46/10090 | | | |
| Gm43318 (ENSMUSG00000105974) | TG vs nTG | 1.918 | 3.34E-02 | | | | |
| Gm43267 (ENSMUSG00000104662) | TG vs nTG | -1.062 | 3.43E-02 | | | | |
| Gm43426 (ENSMUSG00000106214) | TG vs nTG | 1.105 | 3.48E-02 | | | | |
| RP24-96N12.2 (ENSMUSG00000110322) | TG vs nTG<br>TG_HIIT vs nTG_HIIT | -0.335<br>-0.384 | 3.48E-02<br>4.72E-02 | All tissues have low expression<br><br>https://rnacentral.org/rna/URS0002A60790/10090 | | | |
| Gm44386 (ENSMUSG00000107689) | TG_HIIT vs nTG_HIIT | -0.540 | 4.40E-02 | Heart, muscle, lung, adipose tissue, kidney, thymus, skin, colon, trachea<br><br>https://rnacentral.org/rna/URS0002A20769/10090 | ENSG00000233060 | | Brain<br><br>https://rnacentral.org/rna/URS00002EFB52/9606 |
| Arhgap27os2 (ENSMUSG00000085360) | TG_m vs nTG_m | 1.039 | 4.71E-02 | All tissues have low expression<br><br>https://rnacentral.org/rna/URS0002A4F8BB/10090 | ENSG00000294490 | | |
| RP23-299N4.1 (ENSMUSG00000109841) | TG_HIIT vs nTG_HIIT | -0.583 | 4.72E-02 | All tissues have low expression<br><br>https://rnacentral.org/rna/URS0002A4B653/10090 | | | |
| Gm17473 (ENSMUSG00000097805) | TG vs nTG | -0.884 | 4.94E-02 | All tissues have low expression | | | |

| | | | | |
|---|---|---|---|---|
| | | | | https://rnacentral.org/rna/URS0002A49E0F/10090 |
| 9430062P05Rik (ENSMUSG00000104263) | TG vs nTG | -0.467 | 4.95E-02 | |
| Gm17281 (MGP_FVBNJ_G0005910) | TG_HIIT vs nTG_HIIT | -0.337 | 4.99E-02 | |

**Description:**

Sorted by ascending average *p*-adj value across each comparison source.

**Legend:**

TG: transgenic, nTG: non-transgenic, M: male, F: female, HIIT: high intensity interval training, Sed: sedentary, HCM: hypertrophic cardiomyopathy, NF: non-failing, FC: fold change, *p*-adj: Benjamini-Hochberg-adjusted *p*-value
