## Supplementary figures and images for "Investigating the role of long non-coding RNA in hypertrophic cardiomyopathy"

### supp_figs.pdf

Supplemental Fig. S1

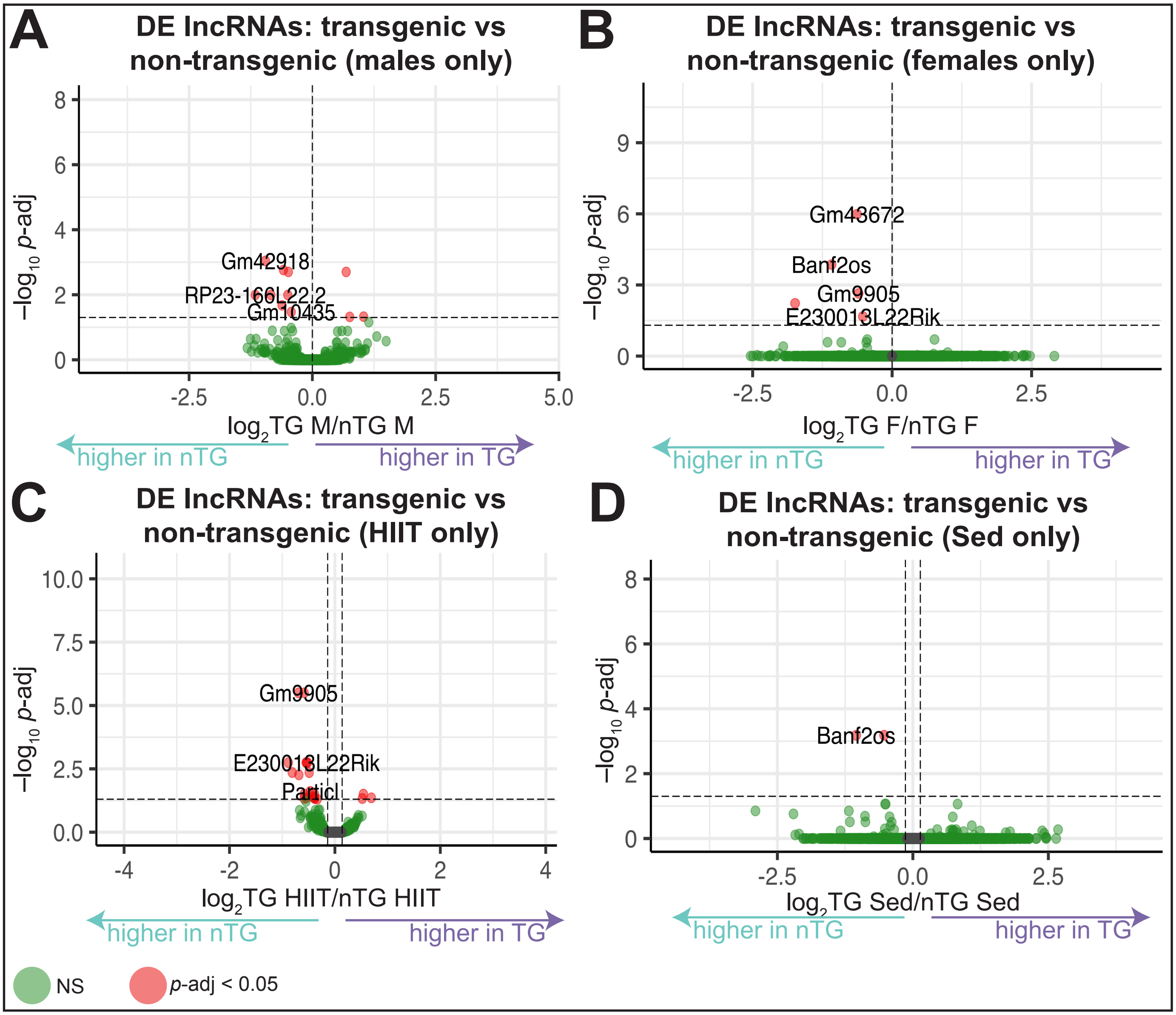

Supplemental Fig. S2

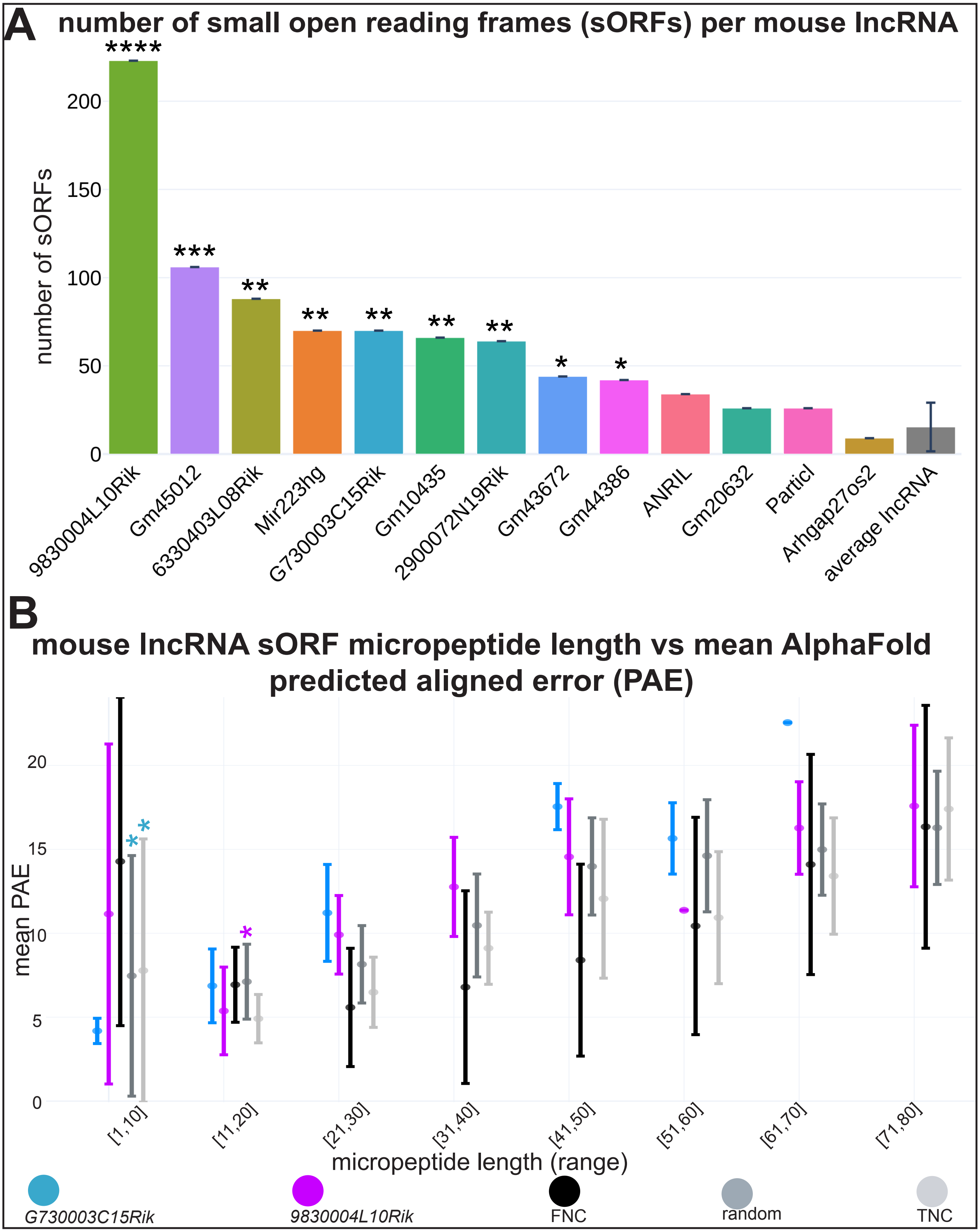

#### Supplemental Fig. S3

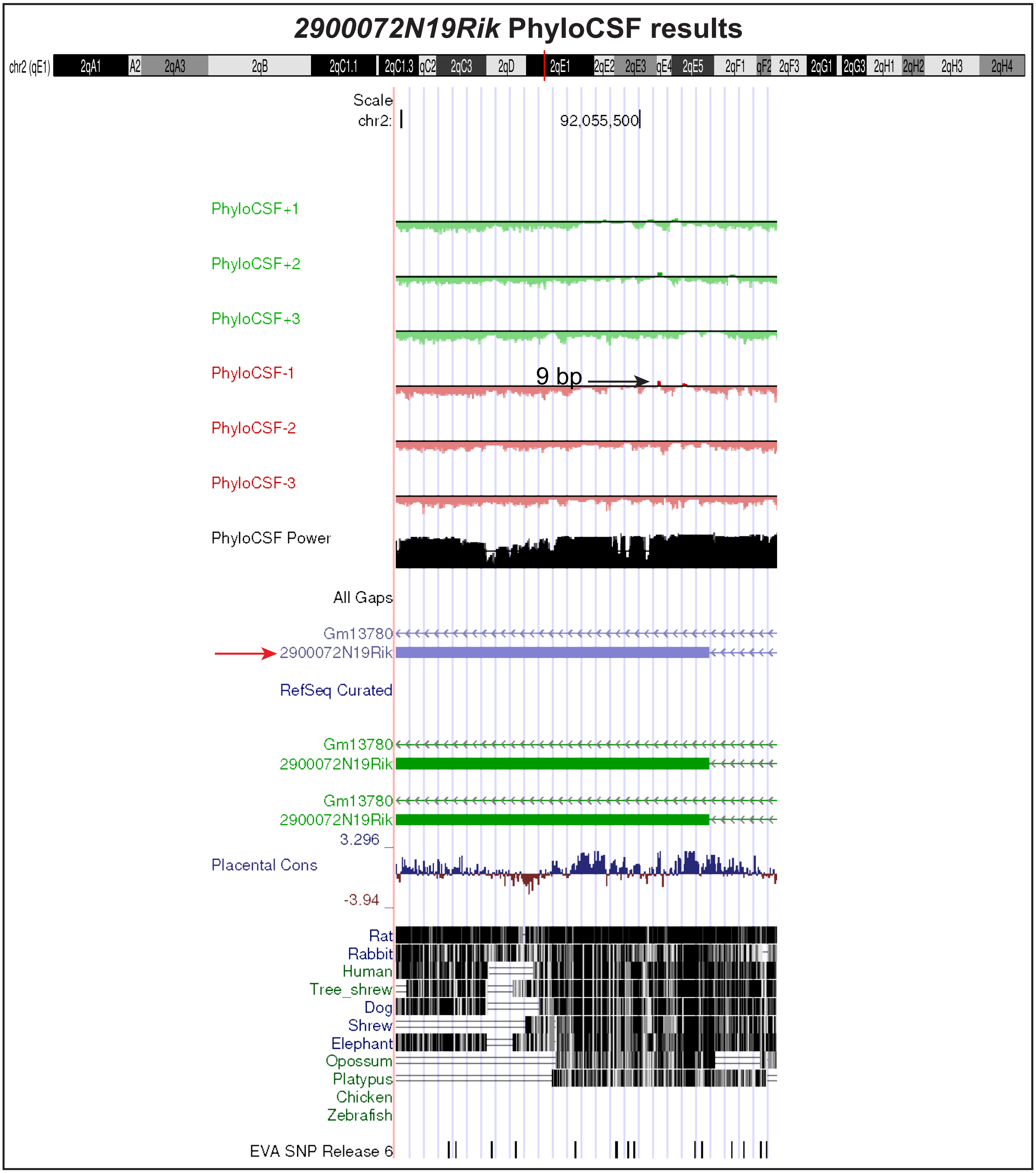
